# Determinants of growth and body size in *Austrolebias* South-American annual killifish

**DOI:** 10.1101/2019.12.31.891648

**Authors:** Andrew Helmstetter, Tom JM Van Dooren

## Abstract

Patterns of size variation in fish are supposed to be generated by growth differences, not by egg or hatchling size variation. However, annual killifish live in temporary ponds with a limited time period available for growth and reproduction. It has therefore been hypothesized that among annual killifish, hatchling size variation should be of large relative importance to generate adaptive adult size variation. Using growth curves of 203 individuals from 18 *Austrolebias* species raised in a common environment, we demonstrate that hatchling size variation indeed is a main determinant of adult size variation in annual killifish, in agreement with the time constraint hypothesis. Furthermore, we find an increased early growth rate in piscivorous species augmenting their difference in size from small congeneric species. This should be adaptive if size differences determine predation success. Environmental effects of spatial location of the population of origin on hatchling size and growth suggest that the time constraint might be weakened in populations occurring near the Atlantic coast. Our study reveals how extreme environments demand specific life history solutions to achieve adaptive size variation and that there might be scope for local adaptations in growth trajectories.

## Introduction

Body size differences are seen as key to understanding life history variation (Roff 1993). Teleost fish alone are spanning nearly nine orders of magnitude in mature size and this is supposed to be due to evolved differences in growth, not to size variation at hatching (Sibly et al 2015). At the same time, there is enormous variation in life cycles among teleost fish and in the ecological and evolutionary variability affecting size differences between closely related species and between and within populations (Hutchings 2002). For example, Atlantic salmon vary 14-fold in size at maturity between populations (Hutchings and Jones 1998) and this variability has been linked to temperature-dependent growth (Jonsson et al 2013).

Size variation and adaptation in fish is much studied in the context of size-selective harvesting (Law 2007). The topic has spurred a modelling effort to support arguments that responses of populations to size-selective harvesting are adaptive (Ernande et al. 2004). Other modelling studies have aimed to predict how competition can affect the emergence of size differences within populations or between species. For example, Persson et al (2000) and Claessen et al (2000) have shown that significant size differences can emerge within fish populations as a consequence of competition for food and cannibalism, leading to so-called dwarfs and giants. Van Dooren et al. (2018) referred to these studies to propose that large piscivorous annual killifish and their prey evolved in sympatry due to a similar scenario of adaptation. Metabolic scaling studies such as Sibly et al (2015) then state that such size differences between fish species must be due to slower or faster juvenile growth, whereas all individuals within a species should grow as fast as environmental conditions for development and metabolism permit. This theory implies that size differences within and between populations can then only be explained by different environmental conditions which individuals experience, leaving little or no room for adaptive variation in the use of resources to achieve particular body sizes. On top of that, expected relative growth rates in fish are expected to decrease with age (Pauly 1979) such that initial growth differences and
 initial environments have larger effects on adult size.

Within the toothcarps (Cyprinodontiformes), annual killifish have evolved at least five times from non-annual species (Furness et al 2015, Helmstetter et al 2016). Annualism denotes that these species inhabit ephemeral waters and that their life history strategy resembles that of annual plants: they establish egg banks where embryos survive dry periods by going through one or several diapauses during development (Wourms 1972). Large size differences among closely related annual killifish species have evolved repeatedly (Costa 2011). Within the genus *Austrolebias* which occurs in South-American temperate environments, large size evolved from small (Van Dooren et al 2018, Helmstetter et al. 2018) and adult body lengths range from about three centimeter to fifteen (Costa 2006), corresponding to a more than hundredfold difference in volume. One of the clades with large species in *Austrolebias* have become specialized piscivores (Costa 2011, Van Dooren et al 2018). In the African genus *Nothobranchius* there is similar size variation involving the evolution of piscivory (Costa 2011, Costa 2018). This genus is well-known for its explosive growth and extremely early maturity, observed both in the lab and the field (Blažek et al 2013, Vrtilek et al 2018). Within killifish, adult body sizes are not only determined by growth variability as Sibly et al (2015) predicted. Eckerström‐Liedholm et al. (2017) found that egg sizes in annual fish are larger than in non-annual toothcarp species and explained this as an adaptation to environments with time constraints on growth periods such as the temporary ponds annual fish inhabit. By being born from larger eggs, annual killifish achieve large (adaptive) sizes by increasing hatchling size instead of growing longer.

We investigated in a common garden lab context and using the South-American annual killifish genus *Austrolebias* how both small and large species in this genus achieve the size differences known from the field and the lab (Figure one). We aimed to identify the major axis among different components contributing to size variation (Schluter 1996): either growth variation (Sibly et al 2015) or hatchling size variation (Eckerström‐Liedholm et al 2017) might explain body size differences more. Hatchlings were obtained from a range of eighteen species mostly occurring in regions close to the Atlantic Ocean and these were raised individually in separate tanks to provide individual growth data. Sizes were measured repeatedly over an eight weeks period. We investigated effects of different environmental variables on hatchling size and growth and compared the patterns of growth rates between the geographic locations of the sites of origin of the populations in our study. We find that hatchling size variation makes the largest contribution to size variation between species, but the relative importance of early post-hatching growth on individual size variation is comparable with hatchling size. Our results thus confirm that large body sizes in some species are to a large extent determined by large hatchling size and we reject the hypothesis that only growth variation matters for size differences between fish species.

**Figure 1.**
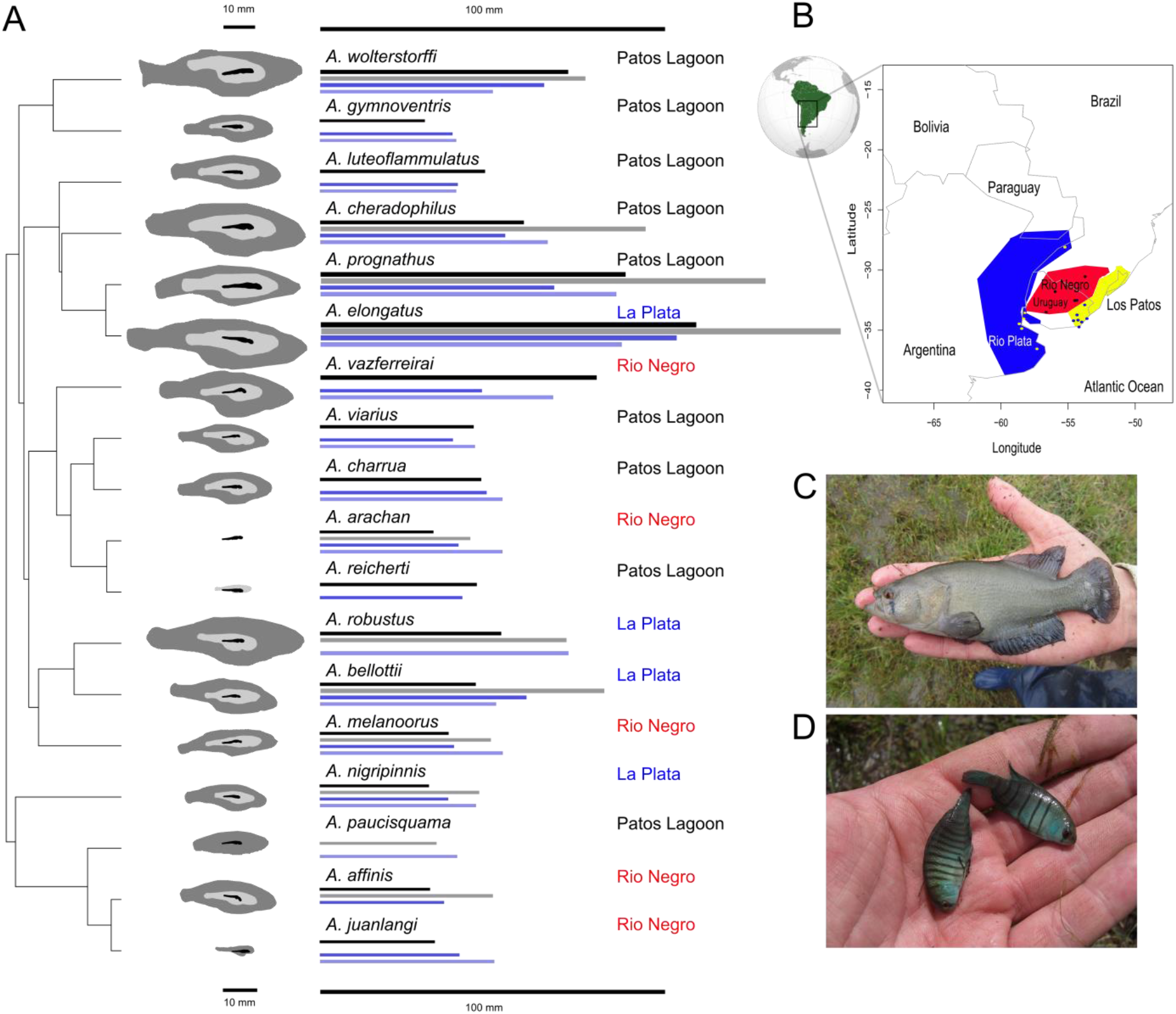
Overview of size data on *Austrolebias* annual killifish from different studies. Per species, silhouettes show the average contour shapes of the species in this experiment at hatching (black), at day 28 (light grey) and at the end of the experiment (dark gray). Bars to the right of the contours indicate standard length data from up to four datasets. Uppermost bar: the size PC used by Van Dooren et al (2018), second bar: maximum sizes used in Helmstetter et al (2018). Third bar: lengths from a lab experiment in Leiden in the Netherlands in 2008. Fourth bar: data collected in 2013 from outdoor breeding stocks at the Foljuif field station foljuif.ens.fr and outdoor breeding in a private garden in the Netherlands. An inset (B) shows the three areas of endemism species in this study originate from. Locations where fish populations originate from are added as points. Inset photos: (C) *A. elongatus* (Photo credit Marcos Waldbillig), which is the largest known *A. elongatus* male; (D) *A. reicherti* (“Paso del Dragon”).

## Material and methods

We triggered hatching of embryos of 18 *Austrolebias* species (Fig. 1) stored in brown peat, by flooding the peat and eggs with water (15 C, 20% aged tap water, 80% RO water, peat extract). Six of the species are usually classified as large and they belong to three different clades (Van Dooren et al 2018, Helmstetter et al 2018). Three species in this experiment are from a single clade of large species containing specialized piscivores (Costa 2010). The 18 species originate from three different Atlantic coastal areas of endemism (Costa 2009), with the populations in this experiment originating from a single such area per species (La Plata river basin, Negro river basin, Patos coastal lagoons, Helmstetter et al. 2018). In these regions, temporary ponds dry in summer. The inland seasonal pattern of rainfall in the Chaco region is different. At the moment of hatching, embryos were between four and forty-two months old.

Hatchlings that were swimming freely (with inflated swim bladder) were placed in separate 0.25 L plastic raising tanks and gradually moved into increasingly larger tanks as they grew. Water parameters were controlled to the following values: <12 dGH, <10 mg/L NO3, < 0.1 mg/L NO2, < 0.25 mg/L NH3, pH = 7.0 - 8.0, 22 ± 0.5 C, by diluting water in the raising tanks daily with water from reserves stored in the same room. The fish experienced a 14L:10D photoperiod. Hatchlings were fed *Artemia salina* nauplii daily for two weeks and then a combination of *Artemia salina*, Chironomid larvae, Tubifex and *Daphnia pulex.* We ensured that the raising tanks always contained live food, such that the fish could feed to satiation. Each tank contained plants (*Vesicularia dubyana* and *Egeria densa*) as well as 5 g of boiled brown peat to aggregate waste and maintain water parameters. At day 58, 32 fish showed visible evidence of stunting or hampered growth (bent spine - extreme lack of growth) and they were assigned to a separate "stunted" category for analysis.

### Photography

We photographed individual fish using a digital USB microscope at hatching (day 1) and repeatedly after that, after intervals of increasing duration with age. We obtained up to nine measurements per individual fish. We constrained the fish in small chambers and photographed from a lateral and dorsal perspective or placed larger fish in a shallow water layer in a petri dish to make lateral pictures only. We measured total length, the distance from anterior tip of the maxilla to the posterior tip of the caudal fin, using ImageJ.

### Statistical analysis

Van Dooren et al (2018) and Helmstetter et al (2018) investigated whether shifts in selection regimes occurred for size and niche traits within *Austrolebias*. Helmstetter et al (2018) identified a shift to a selection regime with an increased optimum size for the clade containing *A. elongatus*. An analysis for shifts on the posterior distributions of phylogenies in Van Dooren et al (2018) found similar shifts for the other clades containing large species in a fraction of trees. Niche traits showed weak evidence for a niche shift in the Negro area (Helmstetter et al 2018). These results make it necessary to use membership of the clades with large species and the areas of endemism as explanatory variables to accommodate effects of the detected regime shifts and to accommodate other similar potential shifts for the traits we investigate. The remaining species differences are then random with respect to these estimated shifts and the species effects can be treated as random effects.

#### Survival and stunting

Next to species differences in mortality, the incidence of stunted body morphologies can indicate whether the environmental conditions we imposed permit normal growth. We therefore assessed survival variation between species and whether the risk of becoming stunted differed between species or depended on age of the embryos at inundation. The proportion of individuals alive at day 50 (before some *A. wolterstorfii* were moved out of the experiment into bigger tanks) was analysed using a binomial generalized additive (GAM, Wood 2017) or generalized linear model (GLM cCullagh and Nelder 1989). Age at hatching, membership of a clade of large species (yielding three categories of large species and one small), area of endemism per species (three areas) and spatial coordinates of the location where the individuals were sampled that the hatchling descended from were used as explanatory variables.

We used scores of the two components of a principal component PC analysis carried out on latitude and longitude of all ponds. The ponds are not randomly distributed across the South-American continent and we wanted to use two independent explanatory variables characterizing spatial locations. The scores were standardized across all observations, so that their averages would be zero across each dataset analysed. We first added scores as thin plate regression splines, hence GAM were fitted (Wood 2017). When model comparisons revealed that these effects should not be retained in the model or when they could all be fitted as linear effects, GLM’s were fitted. Model selection occurred by model simplification using likelihood ratio tests (LRT smooth terms and interaction terms first if present) for comparisons. The probability to become stunted in the experiment was analyzed similarly. We did not include sex effects (male/female/unknown) as an explanatory variable here, as an individual might end up in the “unknown” category due to stunting or premature death.

#### Initial size

We investigated variables affecting initial size at hatching using phylogenetic linear mixed models (de Villemereuil and Nakagawa 2014) and linear mixed models (Pinheiro and Bates 2000). In this manner, unbalanced data can be analyzed while different sources of variation in the data are addressed simultaneously. In each model, we fitted different explanatory (fixed) variables, i.e., embryo age, being classified as stunted, sex, areas of endemism, clades with large species and, as above, we fitted models with the coordinate scores of capture locations. Different random effects were included. In the maximal models, a random species effect with species covariances calculated according the expected values under a Brownian motion model of evolution and next to that a species effect with zero covariances. The expected covariances of the phylogenetic random effects were calculated on the basis of a consensus nDNA tree from Helmstetter et al. (2018). We tested whether including the phylogenetically structured random species effects contributed significantly using a likelihood ratio test. We carried out model selection on the fixed effects as above using likelihood ratio tests (Bolker et al 2009). We used function lmekin() to fit the mixed models including phylogenetic random effects in R (Therneau 2012). For models which did not account for phylogenetic relatedness, we used linear mixed models or generalized additive mixed models (GAMM) with smooth functions to fit the scores of spatial coordinates.

#### Growth

We inspected growth curves y(*t*) of age *t* (days since hatching) using smooth functions (Wood 2017), where we used thin plate regression splines of *t* per species and a smoothness parameter shared between species (factor smooth interaction, function gam from library mgcv(), option “bs=fs”, Wood 2017). We found recent field data on individual size at age for three *Austrolebias* species (Garcia et al 2018) and added these to a figure to check that our common garden environment allowed individuals to grow to sizes comparable to field conditions. From two lab studies, average sizes at different ages were extracted. Errea and Danulat (2001) provided an average total length at age for *Austrolebias viarius* kept in the lab at 25C and Fonseca et al (2013) provide average standard lengths for *Austrolebias wolterstorffi* kept at 24C.

Per individual, we calculated estimates of relative growth per day *g*_*i*_ from the data as

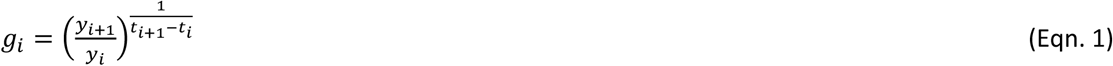

where *y*_*i*_ and *y*_*i* + 1_ are successive size measurements on the same individual at ages *t*_*i*_ and *t*_*i*_ + 1. The standard calculation of relative growth rate is the logarithm of this quantity (Hoffmann and Poorter 2002). We decided to integrate the instantaneous rate over a time interval of one day such that *g*_*i*_ becomes the relative increase per day which is easier to interpret. 100*(*g*_*i*_ - 1) is the percentage relative increase per day. We inspect and analyze relative increases per day at interval midpoints *x*_i_ = (*t*_*i*_ + *t*_*i*+1_)/2. For illustration, we fitted smooth functions to the relative growth *g*_*i*_ evaluated at interval midpoints *x*_*i*_.

We investigated which variables affect relative growth as above, using generalized additive and phylogenetic or non-phylogenetic mixed models. As each analysis contains several measures of relative growth per individual, we added individual random effects nested within the non-phylogenetic species effect. We derived an expression for the error of the relative growth calculations (Supplement), which we implemented in the mixed models. However, it was systematically outperformed by a variance regression for the residual variance which changed with the number of days after hatching (using varExp() weights in lme(), see Pinheiro and Bates 2000).

#### Relative contributions of initial size and relative growth to final size variation

Final size at the end of the experiment (day 58 after hatching) depends on initial size at hatching and on the cumulative growth over the time interval. We can partition cumulative growth into the contributions of different periods to assess their relative importance in the generation of size variation. Log-transformed final size in the experiment consists of additive effects of initial size and growth as shown below. This allows a useful variance decomposition (Rees et al 2010) of final size in the experiment in terms of initial size and growth in different time intervals.

Because day 57 was the last day where the majority of fish were measured, we decomposed final size of an individual *y*_57_ at day 57 of the experiment as follows:

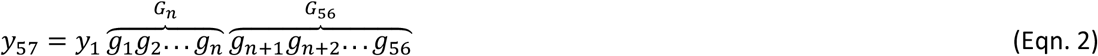

We chose to partition the relative growth per day *g*_*i*_ into two periods, from day one to *n* and from day *n* + 1 to day 56. For the analysis presented, we chose *n* = 28 because this provided two intervals of similar length, with a large number of individuals measured at the end of the first interval. When we log-transform (Eqn. 2), we obtain a sum of contributions to log final size: ln *y*_57_ = *ln y*_1_ + *ln G*_28_ + *ln G*_56_. By means of a variance decomposition of ln *y*_57_ in the variances and covariances of these three terms, we can assess the contribution of each term to final size variation in the experiment (Rees et al. 2010).

For individuals that were not measured on days 29 and 57 but just before or after (14/111 and 14/94, respectively), we extrapolated their sizes to these days (one or two days away), using the relative growth over the last interval in the period. Magnitudes of the three contributions to final size in the experiment were compared with paired samples Wilcoxon tests (Wilcoxon 1945).

The log size variance on day 57 of the experiment depends on the variances of the three components contributing and their covariances (Eqn. 3).

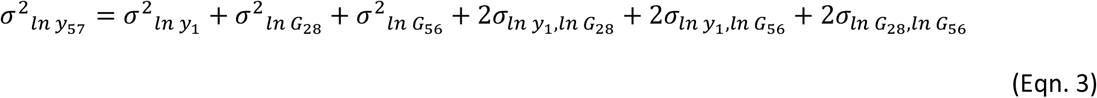

This is the covariance of final size with itself, which is the sum of the covariance of final size with initial size, the covariance of final size with log relative growth until day 29 (early growth), and with log relative growth between days 28 and 57 (late growth). We can interpret absolute values of these three quantities divided by their sum as relative importances (Rees et al. 2010).

We determined relative importances of the three components for the variance between individuals, restricted to individuals that were not stunted and which provided values for all three terms. We resampled the dataset 100 times to obtain standard deviations on the relative importances.

We also calculated relative importances for the variance in final size between species. To obtain estimates of component variances and covariances, we fitted multivariate mixed models (Bates et al 2014) to the three log components of final size with random species effects, allowing random effect covariances between the three component traits and including the fixed effects listed above with component-specific values. In agreement with the analyses of initial size and relative growth rates, we present results assuming random effects per trait which are independent between species. The covariances between the final sizes per species and each size component were then calculated as follows. Predicted values of the summed fixed and random effect part of the mixed model were generated for all species (twelve) where predictions for all three components could be made, To remove individual variation in the fixed effects, we assumed for the predicted values that fish were hatched after four months in the egg, did not show stunted growth and were sexed as a male. The species-specific areas of endemism and taxa with large species were kept as fixed effects. The variances and covariances of the predicted species values were calculated and a parametric bootstrap of the mixed model was used to obtain standard deviations on the relative importances. Magnitudes of the three predicted species contributions to final size were compared with paired samples Wilcoxon tests (Wilcoxon 1945).

#### Comparison with other toothcarps

We compare our results with growth data from other studies on annual and non-annual killifish. We found group averages of size at age, from which we calculated relative growth per day as above. We point out that such estimates based on averages can be biased (Hoffmann and Poorter 2002). We did not observe large changes in variances between pairs of data points from which we calculated growth, therefore we expect such bias to be limited. We retrieved data from studies on *Austrolebias*, *Nothobranchius* and non-annual rivulids and present relative growth estimates we found or calculated. We have added relative growth on one Profundulid for comparison, *Fundulus heteroclitus*, which is a non-annual killifish and a model organism (Schartl 2014). The data file is available as supplementary information. We present a graphical comparison of the results from our experiment with the values obtained from these studies.

## Results

We hatched 203 fish that could swim freely, of which 116 reached day 50 after hatching. The principal component analysis on the coordinates of pond locations resulted in a first PC parallel with the Atlantic coast in a North-South direction and a second PC orthogonal to that, which therefore captures differences in the distance from the Atlantic Ocean.

### Survival and stunting

When we plotted a log survivorship curve of all survival data, we noted that the overall death rate is constant. We found no significant effects of age of the embryos on survival probability until day 50. Species from the (*A*. *robustus*, *A*. *vazferreirai*) clade of large species have a reduced survival probability (*β* = −2.06 (0.70), *χ*^2^(1) = 10.50, p = 0.0012). At the same time, there is an effect of the areas of endemism (*χ*^2^(2) = 9.81, p = 0.007). Species from the La Plata area of endemism have a larger survival probability (estimate difference *β* = 1.37 (0.50), Patos *β* = 0.45 (0.45)). We found a significant effect of area of endemism on the probability to become stunted (*χ*^2^(2) = 16.26, p = 0.0002). There are no stunted individuals among species from the Negro area and about 20% in species from the other two areas. Large species from the *A*. *elongatus*, *A*. *prognathus*, *A*. *cheradophilus* group had an increased probability to become stunted (*β* = 1.57 (s.e. 0.43), *χ*^2^(1) = 13.74, p = 0.0002). Smooth functions of spatial coordinate scores had no significant effects on survival nor stunting.

### Initial size

In GAMM models with all fixed and a random species effect, PC’s of spatial coordinates were best fitted with linear functions. We therefore fitted phylogenetic mixed models with such linear functions to find that the phylogenetic random effect could be removed (LRT non-significant, AIC smaller without phylogenetic covariances, Akaike 1974). From model selection of the fixed effects, we found that the three taxa with large species systematically have larger hatchling sizes. Embryos born from older eggs are larger (Table 1), demonstrating scope for cohort effects and selection on size in the egg bank. Hatchlings of small species from the Patos area of endemism are larger relative to the Negro area, and those from La Plata smaller. The first PC score, which increases in a direction parallel to the Atlantic coast and to the North (called “North” from here on) does not have a significant effect on hatchling size. The second PC increases towards the Atlantic Coast (Called “Coast” from here on) and has a negative effect, hence hatchling size decreases for population situated closer to the Atlantic Coast (Table 1).

**Table 1.** Contributions of explanatory variables to hatchling size and relative growth variation. Parameter estimates and their s.d. for fixed effects of the mixed effect models. Most model parameters are differences from the estimated intercept, which predicts the value of an individual female of a small species originating from the Negro area of endemism. Chi-squared values and tail probabilities of likelihood ratio tests are added when significant for that explanatory variable. “NS” indicates effects that were not significant and removed during model selection.

| Explanatory Variables | Response Variables |  |  |
| --- | --- | --- | --- |
|  | Initial Size | Early relative growth | Late relative growth |
| Intercept (Female, Negro area) | 4.85 (0.24) | 1.045 (0.003) | 1.037 (0.002) <sup>§</sup> |
| Days since hatching | NA | NS | -0.0007 (0.00003) <sup>§</sup><br>$\chi^2(1) = 365.82; p < 0.0001$ |
| Embryo age | 0.016 (0.003)<br>$\chi^2(1) = 21.6; p < 0.0001$ | NS | NS |
| Area of endemism | La Plata -0.37 (0.28)<br>Patos 1.30 (0.33)<br>$\chi^2(2) = 21.2; p < 0.0001$ | NS | La Plata 0.0072 (0.0021)<br>Patos 0.0047 (0.0020)<br>$\chi^2(2) = 10.2; p = 0.0062$ |
| Large clade 1<br>(*A. wolterstorffi*) | 3.09 (0.41)<br>$\chi^2(1) = 23.8; p < 0.0001$ | NS | NS |
| Large clade 2<br>(*A. robustus, vazferreirai*) | 1.70 (0.33)<br>$\chi^2(1) = 17.1; p < 0.0001$ | NS | 0.0073 (0.0030)<br>$\chi^2(1) = 5.17; p = 0.017$ |
| Large clade 3<br>(*A. elongatus, cheradophilus, prognathus*) | 3.96 (0.26)<br>$\chi^2(1) = 44.1; p < 0.0001$ | 0.015 (0.004)<br>$\chi^2(1) = 8.70; p = 0.0032$ | -0.0043 (0.0017)<br>$\chi^2(1) = 5.09; p = 0.024$ |
| Sex | NS | Male 0.0014 (0.0028)<br>Unknown -0.0053 (0.0029)<br>$\chi^2(2) = 6.29; p = 0.043$ | NS |
| Stunted | NS | NS | -0.0084 (0.0011)<br>$\chi^2(1) = 55.89; p < 0.0001$ |
| North | NS | NS | NS |
| Coast | -0.45 (0.12)<br>$\chi^2(1) = 14.1; p = 0.0002$ | NS | NS |

<sup>§</sup>Days since hatching are rescaled, such that the intercept estimates relative growth at day 15.

### Relative growth

Figure 2 shows growth curves for all individuals in the experiment, and fitted smooth growth curve functions. The figure shows that relative to other studies and to field data, individuals in our lab environment have similar or larger sizes for their ages. Moreover, in our experiment fish seem slightly larger than in the field for their age. The growth data we collected is therefore relevant. Moreover, individuals growing somewhat slower in our experiment are still achieving sizes comparable to individuals in the field. Figure 3 shows the pattern of relative growth across species. Most species initially increase in total length by about 5% per day and by the end of the experiment, they still do so by about 2% per day on average. Fig. 3 shows that there is much more individual variation around the species-specific averages for the first days after hatching. We therefore analyzed daily relative growth until day 15 after hatching and after day 15 separately, thus separating the dataset into two subsets with comparable numbers of intervals per individual.

**Figure 2.**
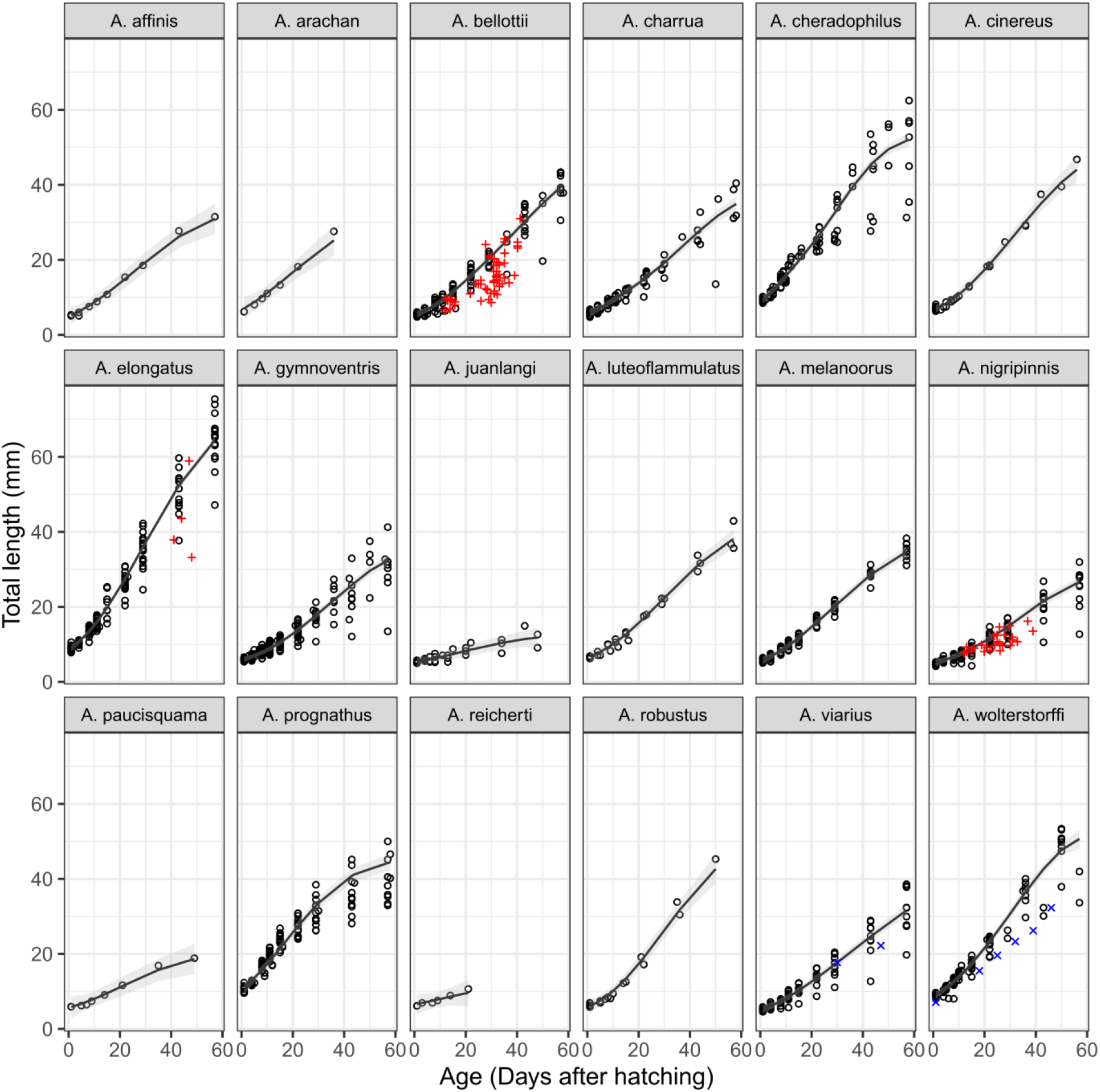
Overview of growth curves of the different *Austrolebias* species in our dataset. Age is expressed as number of days after hatching. Per species, the growth curve predicted by a smoothing spline with a smoothness parameter shared by all species is added. Only the data on non-stunted individuals were used to fit smoothing splines. For comparison, data points from other studies on some of the species we measured are added and colour-coded as follows. Red: individual size-at-age data in the natural environment. *Austrolebias bellottii*, *A. nigripinnis* and *A. elongatus*: individual total lengths at age from the Garcia et al. 2017 field study. Blue: Average size at age. *Austrolebias viarius*: total length lab data were taken from Errea and Danulat (2001), *A. wolterstorffi* standard length lab data from Fonseca et al (2013).

**Figure 3.**
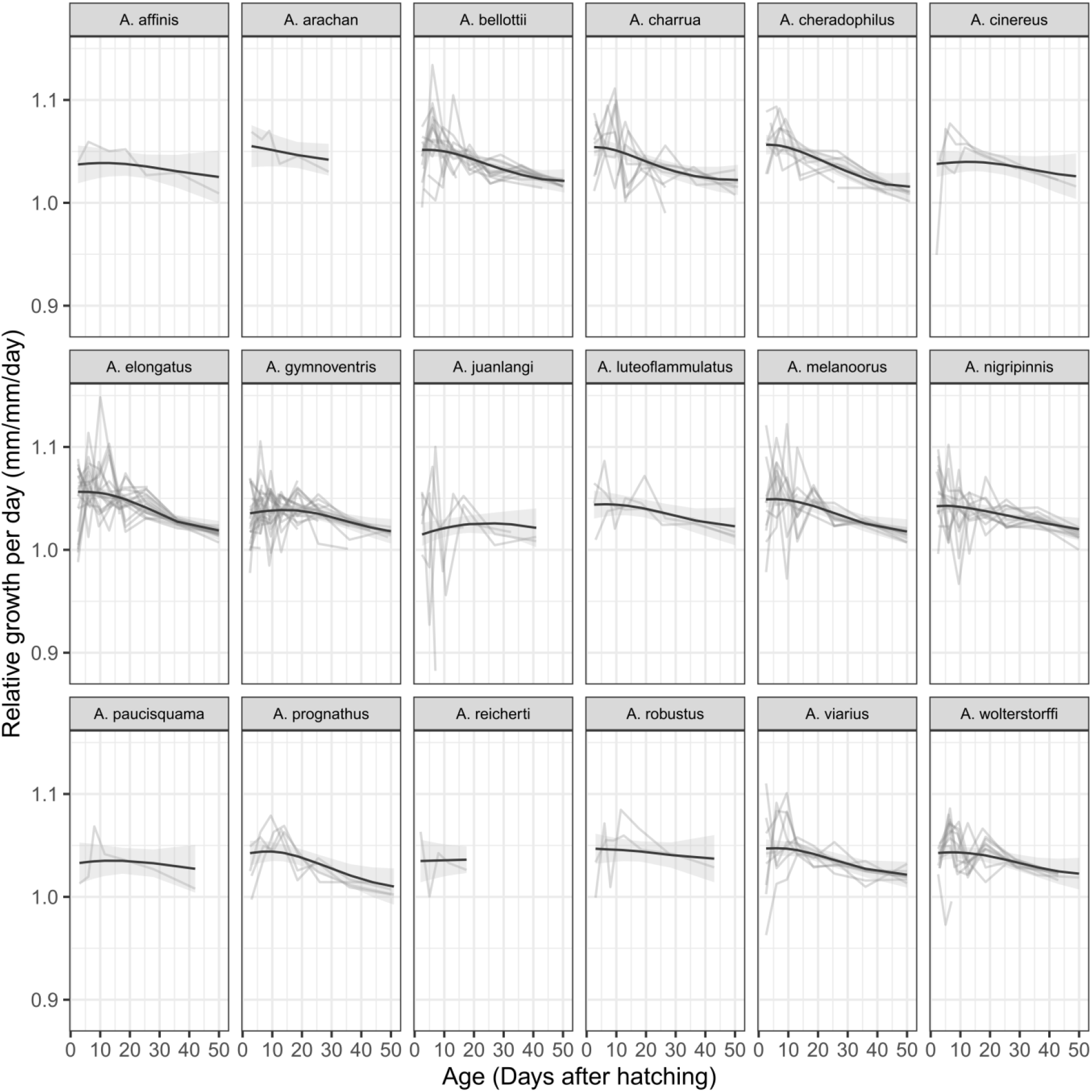
Relative growth per day for the different *Austrolebias* species in this study. Grey lines indicate individual growth histories. Black lines show fitted smooth functions with confidence bands added as in figure two. Data on stunted individuals are not shown.

Different factors affect species differences in different stages of growth (Table 1). Regarding growth during the first fifteen days, a model with phylogenetic random effects did not outperform a model with independent species effects (AIC −2495 vs. −2497, no difference in log-likelihood of the fitted models). Table 1 thus presents a model with independent species effects. Species from the clade containing *Austrolebias elongatu*s grow more rapidly than the other species, about 1-2 % faster per day. Individuals that could not be sexed by the end of the experiment were growing slower shortly after hatching (Table 1). When splines of the PC’s of spatial locations were fitted, these contributed significantly in the complete model, but did not do so after model selection.

Relative growth later in the experiment is still above 3% per day but declines to below one percent per day at the end of the experiment. Again, a model including phylogenetic next to independent species effects was not preferred and Table 1 presents results from the model with independent species effects only. Species from the *Austrolebias elongatus* clade grew slower, whereas *A. robustus* and *A. vazferreirai* grew faster. Stunted individuals grow slower. Species from the La Plata and Patos assemblages grow faster per day, with the largest effect for the La Plata species. When we added spatial coordinates in a GAMM, linear functions of them performed best but these were not retained after model selection. Note that we did not detect any significant sex-specific effects on growth.

### Contributions to final size variation

When we inspect the three log-transformed components of final size (Figure 4), hatchling size clearly makes the largest contribution to final size in the experiment. The contribution of initial size to log final size is significantly larger than that of early growth. The early growth contributions are larger than late growth (both paired Wilcoxon tests *p* < 0.0001, Figure 4). Across individuals, initial size contributes 0.65 (s.d. 0.10) in relative importance of the final size variance, early growth 0.35 (0.11) and growth after day 28 contributes 0.003 (0.060). Initial size thus has a significantly larger relative importance than growth in the second month after hatching. The last component has a small relative importance because the large negative covariance between initial size and growth after two weeks cancels the variance of late growth. When we compare species averages in the figure, it appears that initial size explains most of final size variation among species, paired Wilcoxon tests comparing magnitudes are significant (*p* = 0.0005). There is again a negative covariance between initial sizes of different species and late growth which is larger than the variance between species in late growth. The relative contribution of initial size to final size variance among species is 0.69 (0.07), of early cumulative growth it is 0.19 (0.09) and growth towards the end of the experiment contributes 0.12 (0.08). The confidence intervals for relative contributions among species do not overlap between initial size and early or late growth.

**Figure 4.**
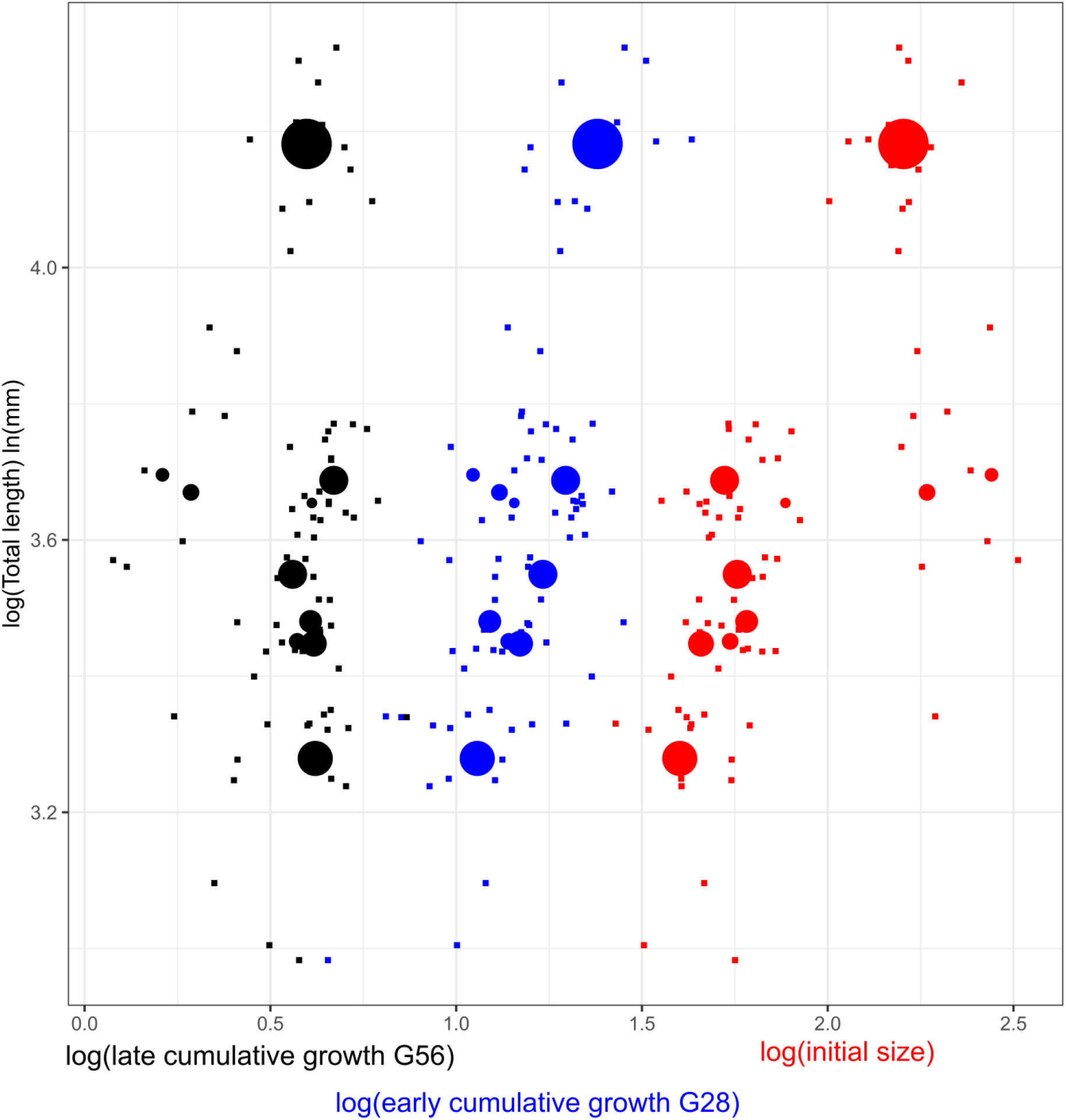
Contributions of log initial size, early and later growth to total size in *Austrolebias*. Individual data points (small squares) are shown for log initial size (red), log cumulative growth from day 1 to 29 (blue) and log cumulative growth from day 29 to 58 (black). Only individuals that were not stunted and that survived until day 56 are included. Per species, average values are shown as circles with the same colours per component as for the individual data. The three top circles are the average components and total size at the end of the experiment for *A. elongatus* (average log of the total length in mm, 4.16), the three bottom circles *A. nigripinnis* (average log total length 3.28).

### Comparison with *Nothobranchius* and non-annuals

When we plot relative growth for all individuals in this dataset (Figure 5) and values from the literature from other related species we see that other estimates for *Austrolebias* are similar to the values we collected. However, in this experiment, individuals sustained levels of relative growth (2-3 %) for much longer. The data on non-annual killifish suggests that these have smaller relative growth rates throughout. *Nothobranchius* fry initially indeed grow explosively, but drop to relative growth rates below the ones in this experiment after three weeks. We note that relative growth in the first weeks for *Nothobranchius* is within the range of measurements we made. We can assume that extremely large relative growth rates in our data are due to measurement error. Alternatively, the data could suggest that some individuals in this experiment are not growing much slower than the average *Nothobranchius*.

**Figure 5.**
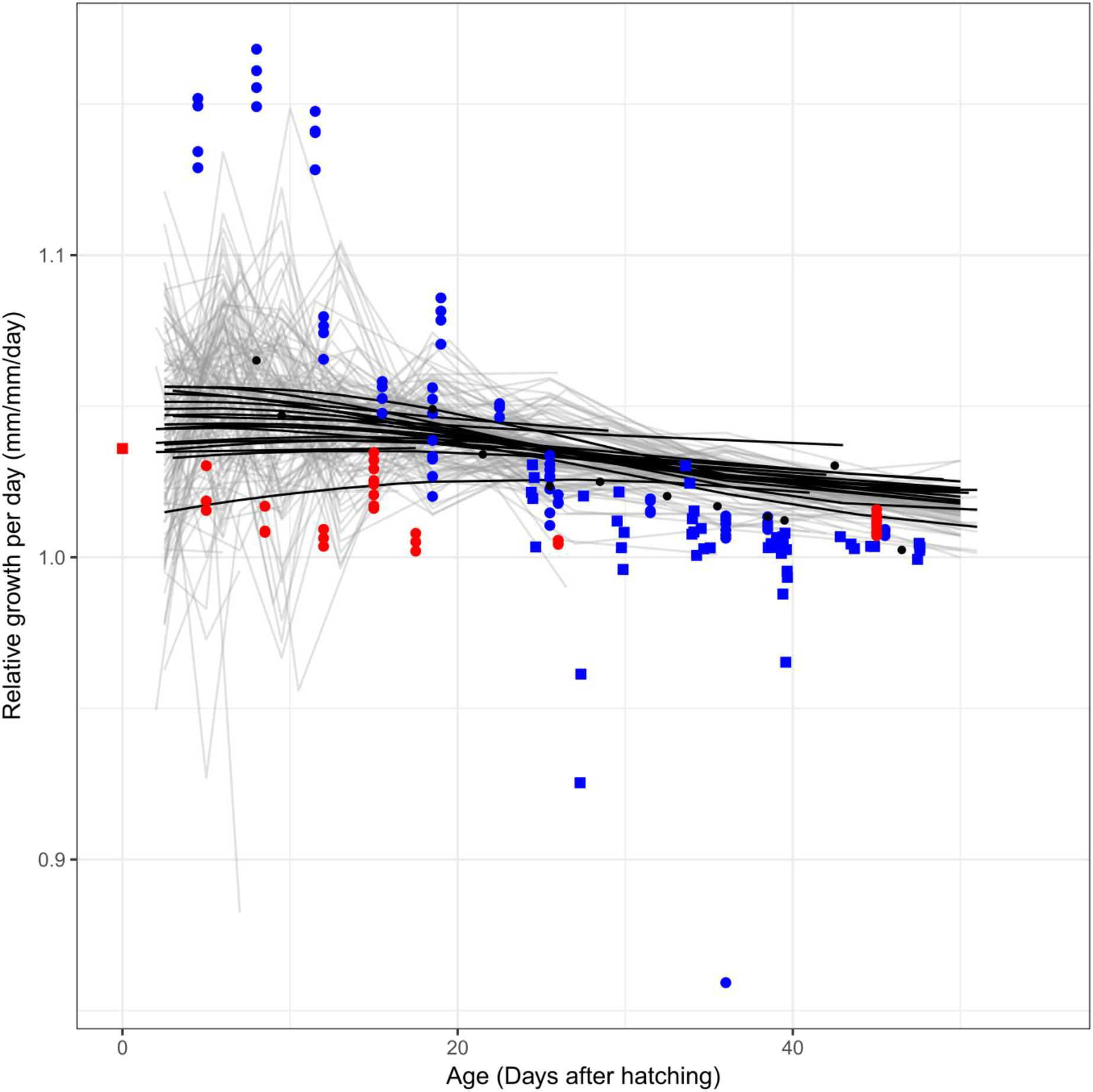
Comparison of relative growth per day in *Austrolebias* with other studies on *Austrolebias* (black points), *Nothobranchius* annuals (blue) and non-annuals (red). The individual field data added in Fig. 2 is omitted here. Other studies did not provide individual values, therefore relative growth was estimated from average sizes at age. *Austrolebias* data of this study are plotted per individual (grey) and the smooth curves from Figure 3 per species are added (black). Squares: field data, circles: lab data. The square at age zero is a relative growth rate estimate for *Rivulus hartii* obtained from field data, but it was unclear at which age the estimate applied.

## Discussion

Hatchling size is the largest contributor to size variation between *Austrolebias* species and its relative importance is significantly larger than that of early or late growth. It is not only determined by species differences, but also by parental or environmental effects, since we found effects of storage duration on hatchling size, of area of endemism and of the distance of the site of origin from the Atlantic coast. Large species from two clades show different patterns of growth over the experiment than smaller species. The *A. elongatus* clade grows faster than the other species in the first two weeks after hatching, but then has a reduced relative growth rate comparable to the smaller congenerics, which we suggest is potentially due to constraints from experimental conditions. The *robustus* group grows faster than the other species from two weeks after hatching until the end of the experiment. This indicates that different clades of large species may be reaching their mature sizes using different growth strategies.

### Adaptive initial size and growth patterns

Individual relative growth rates which are decreasing with age after hatching are adaptive when mortality increases with individual relative growth rate, when mortality decreases with size (Sibly et al 1985). Without environmental changes, catch-up growth is not adaptive (Sibly et al 1985). We observed that the rate of death in our experiment is approximately constant, so at least in the context of our experiment the first explanation does not hold overall. We find, within the experiment, a reduced survival probability for the species of the *A. robustus* clade, and an elevated probability of becoming stunted for the *A. elongatus* group of species. There is therefore no evidence of decreased mortality rates with size, rather the opposite is suggested, but in field conditions the pattern might occur nevertheless. Given that the fish in our experiment grew faster than the available field data, a constraint might be present in the field and affect the adaptive pattern of growth but we do observe some catch-up growth in the *A. robustus* clade of large species, contradicting Sibly et al (1985). The adaptive explanations proposed by Sibly et al (1985) are therefore not supported by the experiment and would depend on field conditions such as competition.

We can also reject the main expectations of Sibly et al (2015): we did not find that all size variation between species is due to changes in juvenile growth. Secondly, within species, there is substantial remaining relative growth variation even when excluding stunted individuals. More specific for the ecology of annual killifish, our results are in agreement with Eckerström-Liedholm et al (2017). We found a large effect of hatchling size variation on final size and all large species have increased hatchling sizes. However, we also found differences in growth among species which contribute to size variation, most notably the increased early growth rate for the largest species. Our finding that hatchling sizes are smaller closer to the Atlantic coast might indicate that individuals are less constrained there by seasonal variation to achieve an adaptive adult size. I.e., near the coast, the seasonality of rainfall might permit longer growth seasons. However, species from the Patos area of endemism which is overall close to the coast initially have larger hatchling size, contradicting this at the between-species level. In addition, we observe that species from the La Plata area of endemism grow faster later after hatching as well as those from the Patos area, to a lesser extent. This might again indicate that there is scope for growth during a longer period after hatching near the Atlantic coast.

We also briefly discuss three additional hypotheses on growth variation. First, predation can select for faster growth. However, we do not know which populations lack predation, except for the Negro area where no piscivorous *Austrolebias* occurs. Second, Arendt (1997) stated that growth can be limited because the rate at which morphological structures develop is limited. For example, muscle structure differs in dependence on growth speed, and can become less efficient with faster growth. The increased growth rate in the piscivorous species after hatching motivates a further investigation to check if these species would sacrifice performance efficiency for size. Third, Dmitriew (2011) explained such costs of growth acceleration in purely ecological terms. When energy allocation is directed elsewhere for example to reduce the time to complete a stage in development, growth must be reduced. It is unclear whether hatchlings of piscivorous species would need to achieve a certain size as soon as possible to permit access to specific resources such as fish prey.

### Comparative lab experiments versus data from the field

Comparative studies such as Eckerström-Liedholm et al (2017) use lab or field data, or both. Size measures from field populations are widely available, but growth rates are often only available as population averages, or rates calculated from size measurements on different groups of individuals (e.g. Winemiller and Rose 1992). An advantage of field data is that it can be assumed that each species has been sampled in an environment it is adapted to. On the other hand, intra- and interspecific competition can affect different species to a different extent, modifying pairwise size comparisons. We have collected lab data for a comparative analysis. With lab data obtained in one or several controlled environments, it is likely that some species will be performing less than others in the chosen environments. Hence, some species will show their overall maximum growth rates while others may not. To understand the causation of size variation, field data don’t seem a valid substitute for controlled lab experiments, but they can be used to assess the pertinence of growth patterns observed in the lab. If the purpose is to compare adaptive growth curves between species, environments tuned to each species or field environments seem required.

Martins and Hansen (1996) pointed out that comparative methods often have the same weaknesses as meta-analyses, and at the time, methods didn’t permit incorporating individual variability easily. In addition, Goolsby (2015) noted that field data might render inference unreliable when it assumes the absence of phenotypic plasticity. With the advent of phylogenetic mixed models and the realization that these models are similar to the animal model of quantitative genetics (Lynch 1991), it has become easier to analyse lab data obtained in complex experimental designs and environments. We propose to see our data as character states sampled on the species and individual-specific reaction norms at a particular combination of environmental parameters.

Future studies could expand on the environmental treatments imposed and will permit to estimate species variation for growth plasticity. We did not need the function-valued methods proposed by Goolsby (2015) to reconstruct ancestral states and maybe infer selection regime shifts, as we had already obtained hypotheses for shifts in traits for some taxa from other studies and could therefore use these as starting points in this study.

A comparative analysis should not require very many species just to overcome limitations of individual data points or limitations of the methods of analysis (Mitov et al 2018). The larger the number of species in an analysis, the less likely that traits are directly comparable between all of them. It therefore seems most obvious to extend the analysis we carried out to an experiment with a similar set of species crossed over several lab environments, to obtain first estimates of species variation in plasticity. However, in quantitative genetics, large and long-term datasets and improved methods have permitted the study of natural selection and phenotypic plasticity in the wild (Charmantier et al 2014). For comparative phylogenetic methods, mixed models applied to multi-species field data might permit similar advances, but to limit the range of species for which detailed data need to be available, and to limit the range of models to be fitted and compared these might require a priori hypotheses to be tested instead of the automated model selection (e.g. Bastide et al. 2018) which is currently common and demands a large set of species to be included.

### Non-annual and African annual killifish

When we compare relative growth rates at different days after hatching between this experiment and other lab and field studies then it can be noted that early relative growth of *Austrolebias* is faster than of non-annuals but slower than of *N. furzeri* in some experiments. Later on, after about a month, the fish in this experiment outperformed nearly all other values we collected. This might be a side effect of our experimental setup, where we avoided competition and degrading environments, or it might be the case that *Austrolebias* sustain fast growth longer and thus achieve larger adult sizes for the same initial size. The amounts of variability we observed between individuals suggest that it might be possible to tweak environments to obtain relative growth rates closer to the ones observed in *Nothobranchius* (Blažek et al 2013). Faster growth might require experimental conditions with fluctuating temperatures (Boltana et al 2017) and there might be species differences in the extent of this effect. Note that we did not tune the environment to specific species and neither did we generate a sequence of environmental conditions to obtain the largest possible growth rates at any age. We chose a standardized common environment where we expected all species from the three areas of endemism to perform relatively well. The fact that we observed no stunting among the species from the Negro area and a smaller survival probability for that area seems to indicate that the environment we chose is not an environment these species are very well adapted to because it led to the strongest expected survival effects on the fish. Species from the *A. robustus* group have a reduced survival probability and a pattern of growth suggesting catch-up growth. This might also be a side effect of the conditions we imposed, where a non-constant environment might lead to overall faster growth and larger size.

### Conclusion

Using growth curves of 18 *Austrolebias* species, we demonstrate that hatchling size variation is a main determinant of adult size variation in annual killifish. In addition, we find an increased early growth rate in the piscivorous species, augmenting their size. Environmental effects of spatial location of the population of origin on hatchling size and growth suggest that the time constraint which explains the importance of hatchling size variation for adult annual fish size might be weakened in populations occurring near the Atlantic coast. This suggests that the manner in which annual killifish defy the overall expectations on determinants of adult fish size, might be locally adapted to environmental constraints.

## Acknowledgements

We thank Tom Smith, Samuel Perret, Beatriz Decencière and Alexis Millot for help with fish care. Armand Leroi and Vincent Savolainen for comments on a previous version of this manuscript.

## Animal Care and Welfare

Fish used in this breeding experiment were maintained and raised at the CEREEP station in Nemours-St. Pierre, France (approval no. B77-431-1).

## Supplement

If measurement error is the same for length measurements at different ages and equal to *σ*^2^_*y*_, we can calculate an approximation to the measurement error in the relative growth rate (Eqn. S1) using a first-order Taylor expansion of total length *y*,

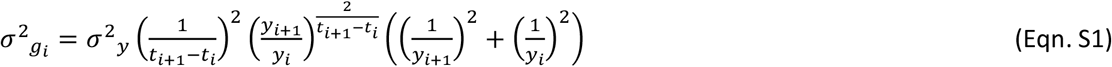

We included this error model in linear mixed models for relative growth rate variation. However, there is no software available to combine such error models with phylogenetic mixed models. We therefore fitted independent species effects (Pinheiro and Bates 2000). As an alternative to this error model, we also allowed the residual variance to depend on the age of the individual. The likelihoods of the data assuming either of these models were compared, also with the likelihood obtained from the model assuming a homoscedastic residual variance. We found that the model where the residual variance depended on individual age outperformed the other two models for early relative growth. We report here the fixed effect tests of that model. For late relative growth, homoscedastic errors were preferred, which is the model in the last column of Table 1.

